# Chromid transforms opportunistic pathogen into beneficial symbiont to raise coral’s thermal tipping point

**DOI:** 10.64898/2026.06.19.733351

**Authors:** Clarence W. H. Sim, Hannah H. C. Luk, Joao P. A. Pereyra, Aaron A.R. Loh, Johnson J. H. Lim, Wai Leong, Jonas C. H. Khaw, Sujatha Srinivas, Isabella Tiaras, Paul C. Kirchberger, Lionel J. W. Lim, Lionel C. S. Ng, Lindsey K. Deignan, Jani T. I. Tanzil, Rebecca J. Case

## Abstract

Resilient turbid coral reefs, found 1° north of the equator, experience fewer and less intense bleaching events despite being situated within the world’s busiest shipping port in highly urbanised Singapore. We hypothesised that bacteria within the coral holobiont play a role in maintaining coral health by conferring traits that enhance thermal tolerance. Eleven *Pseudovibrio* isolates were obtained from the scleractinian coral *Pachyseris speciosa*, and their genomes differed by only four single nucleotide polymorphisms. A ∼490 kbp chromid, pCJH, was present in 7 of the 11 isolates. A *Pseudovibrio* isolate without pCJH is an opportunistic pathogen of *P. speciosa*, accelerating bleaching disease. In contrast, the chromid reduces *Pseudovibrio’*s fecundity, enhances its host colonisation and consequently increases *P. speciosa’s* thermal tolerance by 1 °C. Interaction analyses in experimentally heat-stressed corals revealed a negative interaction between *Pseudovibrio* and *Vibrio* ASVs. The *Pseudovibrio* genomes and chromid encode several diffusible antibiotic biosynthetic gene clusters and two type six secretion systems (T6SS) for contact-dependent inhibition, with both types of inhibitory activity shown against opportunistic coral pathogens from local (i.e. isolated from the same coral host) and type-strain *Vibrio* spp. Together, these findings support the Coral Probiotic Hypothesis that bacteria enhance coral resilience, and identify a chromid responsible for *Pseudovibrio*’s climatically driven Jekyll-and-Hyde lifestyle from opportunistic pathogen to beneficial symbiont.

## Introduction

Within the coral holobiont, bacteria influence the holobiont’s ecology to shape the coral’s physiology and ecological resilience [1, 2]. The Coral Probiotic Hypothesis places their microbe’s ecological interactions and chemical biology within an integrated host system [3, 4]. Consequently, screening to identify beneficial microorganisms for corals (BMCs) has commonly focused on functional traits such as pathogen inhibition, stress mitigation, and the production of beneficial metabolites. The ability of candidate BMCs to establish and persist within the dynamic coral environment is also assessed [5, 6]. Host colonisation is important for BMC efficacy, as stable host association and ecological roles within the holobiont facilitate long-term persistence. Once established, these organisms can mediate numerous processes including metabolic exchange, pathogen suppression, immune modulation, and modification of host-associated biogeochemical cycles [1, 7, 8].

Equatorial coral reefs are ecosystems defined by constant high sea surface temperatures and minimal seasonal variability [9]. At the same time, these reefs exist within one of the world’s most densely urbanised and anthropogenically-impacted coastal regions, characterised by continuously high levels of turbidity and sedimentation that impose multiple stresses on the coral communities [10]. Despite impacts from increasingly frequent and severe marine heatwaves that have driven widespread coral bleaching [9, 11, 12], Singapore’s reefs have persisted, with high taxonomic albeit reduced functional diversity, suggesting long-term ecological adaptation to challenging environmental conditions [13–15]. These conditions have acted as strong environmental filters shaping various aspects of reef ecosystem functioning [16], and may therefore also do so for coral-associated bacteria [4], making equatorial turbid urban reefs promising systems in which to identify taxa with traits that support coral resilience.

The coral holobiont is dynamic and can rapidly reorganise in response to environmental change. Shifts in microbial taxa or functional traits may enhance coral resilience by buffering the effects of environmental stress, whereas dysbiosis can disrupt these interactions and promote the proliferation of opportunistic pathogens [7, 17]. While many reef systems report the widespread dominance of *Endozoicomonas* in healthy corals [18], it is frequently absent or found at overall low abundance in Singapore corals. Where it does occur, however, it can reach high relative abundance and persist even in highly disturbed environments [19]. This suggests that other bacterial symbionts may support holobiont stability in these reefs. Several corals exhibit considerable variation in Symbiodiniaceae populations and bacterial communities across turbid reefs [20, 21], potentially contributing to coral persistence in Singapore’s reef environments [22]. The coexistence of high symbiotic diversity, chronic environmental stress, and persistent coral populations suggests that equatorial urban reefs may harbour BMCs that contribute to coral resilience in the busiest shipping hub in the world.

Despite persistently warm conditions that favour disease through stressing the coral and activating virulence in opportunistic pathogens, coral reefs in Singapore exhibit relatively low mortality from large-scale bleaching compared with many other tropical reef systems [9, 12, 23]. One possible explanation is that BMCs limit the growth or virulence of opportunistic pathogens. *Vibrios*, such as *V. coralliilyticus* and *V. shiloi*, are opportunistic coral pathogens with temperature-dependent virulence [24, 25] that have rarely been isolated from Singaporean corals despite the presence of environmental *Vibrio* populations in surrounding seawater [26, 27]. This raises the question: does the absence of pathogens or the presence of BMCs drive the low incidence in bacterially mediated disease in Singapore’s coral reefs? Moreover, because temperature influences bacterial growth rates, virulence factors, host stress, and gene expression, elevated temperature is a critical environmental parameter for understanding how bacterial interactions within the coral holobiont change as the host becomes thermally stressed.

The bacterial isolates characterised in this study have a brown colony morphology, characteristic of the antibiotic tropodithietic acid (TDA), and were obtained from the coral *Pachyseris speciosa,* a species with relatively high resilience to bleaching that is common on Singapore’s turbid reefs [12, 21]. Bioassays and manipulative experiments accordingly focused on this coral. This study aims to characterise these isolate’s interaction with the coral and with other bacteria, and to identify their genomic and phenotypic potential through genome sequencing, bioassays and manipulative coral holobiont experiments. Critically, we aim to understand how these interactions change under thermal stress. We hypothesize that equatorial turbid reefs contain bacteria that extend the thermal range of corals, by modulating their interactions with other bacteria and the photosymbiont. We therefore:

(1) identified the brown colony forming isolates through genome sequencing;
(2) tested if the *Pseudovibrio*s influence the thermal tolerance of *P. speciosa*;
(3) examined how heat stress alters *Pseudovibrio* interactions with other holobiont bacteria; and
(4) screened the isolates for inhibition of *Vibrio* pathogens.

## Materials and Methods

### Bacterial strains

Bacteria were grown using marine broth (Difco 2216) (MB) at 30 °C for 24 h unless otherwise specified. Liquid cultures were grown at 160 rpm, and agar plates used 1.5% Bacto agar. Two *Vibrio* type strains, *Vibrio coralliilyticus* YB1 [24] and *Vibrio harveyi* 384 (ATCC 14126) [28], were used in this study, in addition to isolates from *P. speciosa* (detailed below). Growth curves were generated by growing liquid cultures in MB at 30 °C, using a Tecan Infinite 200 PRO microplate spectrophotometer. Colony forming units (CFU) were calculated from MB plates. Bacteria were isolated from fragments of *P. speciosa* obtained from the coral fragment library at the St John’s Island National Marine Laboratory (SJINML) as described [29–31].

### Identification of candidate BMCs in corals with different tolerance profiles

To identify potential BMCs that could be targeted for bacterial isolation, four scleractinian coral species, *P. speciosa*, *Goniastrea pectinata*, *Pocillopora acuta*, and *Porites lutea*, were chosen for their varying bleaching tolerance and resilience profiles and high abundance in Singaporean reefs [12, 32]. Fragments of each coral species (n = 6) were collected from Pulau Satumu, Singapore (1.1599, 103.7400) at two time points: early March 2024 (with no apparent bleaching) and late May 2024 (with observed bleaching). Samples collected for molecular analysis were placed in RNAlater and stored at −80 °C.

### Molecular community analysis

16S rDNA metabarcoding was used to characterise the bacterial community associated with coral fragments collected from the field before and during a heat wave in 2024. DNA was extracted from coral tissues using the DNeasy PowerSoil Pro Kit (Qiagen, Hilden, Germany) following the manufacturer’s instructions, with an additional homogenisation step using the FastPrep-24™ 5G Instrument (MP Biomedicals, Singapore) for three cycles at 6 m/s for 30 s, with 30 s on ice between cycles. The V4 region of the bacterial 16S rRNA gene was amplified with primers 515F (5’-GTGYCAGCMGCCGCGGTAA-3’) and 806R (5’-GGACTACNVGGGTWTCTAAT-3’) [33] modified to include Illumina adaptor sequences.

PCR was carried out at 94 °C for 3 min, 94 °C for 45 s, 57 °C for 60 s, and 72 °C for 90 s for 25 cycles, 72 °C for 10 min. Upon purification, sequencing was performed on an Illumina MiSeq platform using the MiSeq Reagent Kit v3 to generate 300 bp paired-end reads at the SCELSE Sequencing Facility, NTU, Singapore. Sequences were processed in R using DADA2 v1.34.0 [34] to generate amplicon sequence variants (ASVs), with additional details in Supplementary Material. ASVs were taxonomically assigned using SILVA v138.2 [35] via *phyloseq* v1.50.0 [36] with a bootstrap confidence threshold of 80/100 and then numbered in decreasing order of relative abundance. ASVs were assessed manually for patchy distribution across healthy and bleached samples in all coral species.

### Genome sequencing, assembly and annotation

Eleven isolates, all forming brown colonies, were isolated from *P. speciosa* and cultured for genome sequencing and downstream analysis (Table S1). Isolation, genomic DNA extraction, DNA library preparation, and sequencing were performed at the SCELSE Sequencing Facility, NTU, Singapore, as described in Loh et al [29]. Annotations of the assembled genomes were carried out using the Prokaryotic Genome Annotation Pipeline (PGAP) v6.7 [37] for submission to the National Center for Biotechnology Information (NCBI) and with Prokka v1.0.9 [38]. Bacterial genomes were identified using the Microbial Genomes Atlas (MiGA) [39] webserver, using the NCBI-Prok function, with the closest homology based on average amino acid identity (AAI).

### Genomic analysis and comparison

To determine the taxonomic identities and genomic relatedness between the isolates and all publicly available genomes of *Pseudovibrio* type strains (Table S2), calculations were performed using both the Orthologous Average Nucleotide Identity Tool v0.93.1 [40] and the Genome-To-Genome Distance Calculator v3.0 [41] to calculate ANI and digital DNA-DNA hybridization (dDDH) scores respectively. For eleven isolates identified as *Pseudovibrio* sp. (SCP18-21, 23-24, 26-30), genomes were visualised and compared using Proksee [42] (to help identify indels between genomes), and functionally characterised using BlastKOALA v3.1 [43] and eggNOG-mapper v3.0.0 [44] using ortholog assignments from the eggNOG 7 database [45]. PlasmidFinder was used to look for plasmids in all genomes [46]. Analysis of single nucleotide polymorphisms (SNPs) was performed using Snippy v4.6.0 [47]. Identification and analysis of secondary metabolite biosynthetic gene clusters was performed using antiSMASH v8.0.4 [48].

Codon usage biases of the chromosome and chromid were determined by calculating relative synonymous codon usage (RSCU) values, and the bias profiles were compared using Pearson correlation. Annotations of protein-coding genes on the *Pseudovibrio* sp. SCP18 chromid (pCJH) from BlastKOALA v3.1, eggNOG-mapper v3.0.0 and Prokka v1.0.9 were compared, manually curated, and genes were assigned to KEGG functional categories. Where annotations conflicted or were ambiguous, protein sequences were searched against the RefSeq database using NCBI BLASTp and examined for conserved domains using the NCBI Conserved Domain Database and InterProScan v6.

Curated genes were then grouped into manually defined functional categories relevant to host colonisation and symbiosis, based on their annotation, genomic context and reported functions in the literature (complete list in Supplementary Data 1).

### Colonisation of coral host

To determine if the carriage of a chromid affected the colonisation of the coral host by the bacterium, colonisation assays were performed following Banin et al [49]. *Pseudovibrio* sp. SCP18 (with chromid, pCJH) and SCP20 (without chromid) were selected among isolates to measure attachment to *P. speciosa* as they have identical chromosome sequences with the presence/absence of pCJH being the only genetic difference between the isolates. Cells were diluted to a target concentration of 10^6^ cells mL^-1^ in Instant Ocean (no exogenous organic carbon added) in a conical flask with a ∼ 4 cm^2^ fragment of *P. speciosa* and a control flask without a coral fragment. CFU was measured at 0, 2, 4, 6 and 8 h and net attachment was calculated from the difference in CFU in the bacterial suspension (i.e. unattached bacteria) relative to the initial inoculum (i.e. starting concentration) [49]. Flasks were incubated at 30 °C, 35 rpm, and in darkness to minimise addition of photosynthetically-derived organic carbon from the coral.

### Pseudovibrios interaction with heat-stressed coral

Experiments to identify the influence of temperature stress on the coral, *P. speciosa*, were performed by manipulating bacterial inoculations of *Pseudovibrios* at the SJINML Biosafety Level-2 aquaria. Six independent coral fragments were used for each treatment: (1) *P. speciosa*, (2) *P. speciosa* inoculated with *Pseudovibrio* sp. SCP18 (with chromid, pCJH), and (3) *P. speciosa* inoculated with *Pseudovibrio* sp. SCP20 (without chromid). These three experimental conditions were subjected to either constant ambient temperature (30 °C) or acute heat stress by ramping 1 °C d^-1^ (30–37 °C over 7 d) and monitored using Single Pulse-Amplitude-Modulated (PAM) fluorometry to identify impairment of PSII as an indicator of bleaching disease. Coral fragments were prepared by inoculating with the bacterial suspension for 8 h. Then, fragments were transferred from flasks in the dark incubator and glued onto plastic blocks attached to a base plate, before being placed into 10 L aquaria for each treatment. No further inoculation was performed. Additional details on PAM measurements and full schematic on aquaria set up (Fig. S1) are in Supplementary Material.

### Impact of heat-stressed corals on bacterial-bacterial interactions

To investigate the bacterial community response to heat stress of the holobiont, the same four species as in the field collection were used (*P. speciosa*, *G. pectinata*, *P. acuta*, and *P. lutea*). Six replicate fragments from each species were sampled from constant temperature (29 °C) and heat-ramping (1 °C d^-1^ increase from 29–37 °C for 8 d). Acute heat stress was used as a controlled bioassay to quantify the temperature-dependent inhibition of photosystem II (PSII) at 1 °C increments, to simulate an ecological heat-wave scenario in a rapid experiment (i.e. to reduce adaptation). Aquaria were set up in 45 L tanks with a 12:12 h (light:dark) diurnal cycle at 120 µmol m^-2^ s^-1^ PAR. Fragments in the heat-ramped aquaria were assessed every day for bleaching using a colour reference chart and sampled when bleaching was first observed visually (i.e. near white in colour): at 34 °C for *P. speciosa* and *G. pectinata*, and 35 °C for *P. acuta* and *P. lutea*. Their corresponding fragments in the control aquaria were simultaneously sampled for molecular analysis.

To identify potential associations between community members, interaction networks were constructed using *SpiecEasi* association algorithm [50] independently for the control and heat-treated samples using 16S metabarcoding as described in subsection “*Molecular community analysis”*. The threshold value for associations detected was set using the default value of 0.1. Multiplicative simple replacement (multRepl) was used for zero replacements in the ASV abundance matrices, which were treated with centered log-ratio (CLR) transformation. BLAST was used to align *Pseudovibrio* and *Vibrio* ASVs with the 16S rRNA gene fragment sequences of the SCP strains. ASVs showing significant interactions with *Pseudovibrio* ASV0135 were extracted and plotted using *ggraph*.

### Inhibition assay

Three *Vibrio* isolates from *P. speciosa* represented three species (*Vibrio* sp. SCP2, *V. coralliilyticus* SCP4, and *V. harveyi* SCP5) as they had ANI < 95% and dDDH < 70% when comparing them to each other, with SCP4 and SCP5 having ANI > 95% and dDDH > 70% to respective type strain genomes. *Vibrio sp.* SCP2 represented a candidate novel species, as it had no matches with ANI > 95% and dDDH > 70% in public databases [27]. To assess the antagonistic activity of *Pseudovibrio* isolates against coral-associated *Vibrio*, isolates were screened for their interaction with Singapore coral isolates SCP2, SCP4, SCP5 as well as *V. coralliilyticus* YB1 and *V. harveyi* 384, which were not isolated in Singapore. Soft agar overlays were made with *Vibrio* SCP cultures, and overnight cultures of *Pseudovibrio* isolates were used for drop-plating. Zones of inhibition were measured after 48 h of incubation and assessed for patterns consistent with contact-dependent or diffusible inhibition.

### Statistical analysis

Statistical analysis of results from growth curves, colonisation assays and measurements from all heat-ramping aquaria experiments were performed using SigmaPlot v16.0. Pairwise comparisons used t-tests, and comparison of multiple groups was performed using analysis of variance (ANOVA) and the Holm-Sidak test. Comparison of growth curves used the Mann-Whitney rank sum test, with exponential growth occurring between the lag and stationary phase and calculated between two time points within this period (between 8 and 12 h for strains without chromid; between 12 and 16 h for strains with chromid).

## Results

### Heterogeneous distribution of Pseudovibrio among coral species

To assess the distribution of *Pseudovibrio* across coral hosts, we examined its relative abundance in the bacterial communities of healthy and bleached colonies of *P. speciosa*, *G. pectinata*, and *P. lutea* using 16S metabarcoding (Fig. 1A). In healthy corals, *Pseudovibrio* was detected in all three coral species, occurring in more individual samples of *G. pectinata* than of the other two species. In bleached corals, the relative abundance of *Pseudovibrio* was lower in *P. speciosa* and higher in *G. pectinata* than in their healthy counterparts, whereas it was not detected in bleached *P. lutea*.

**Figure 1:**
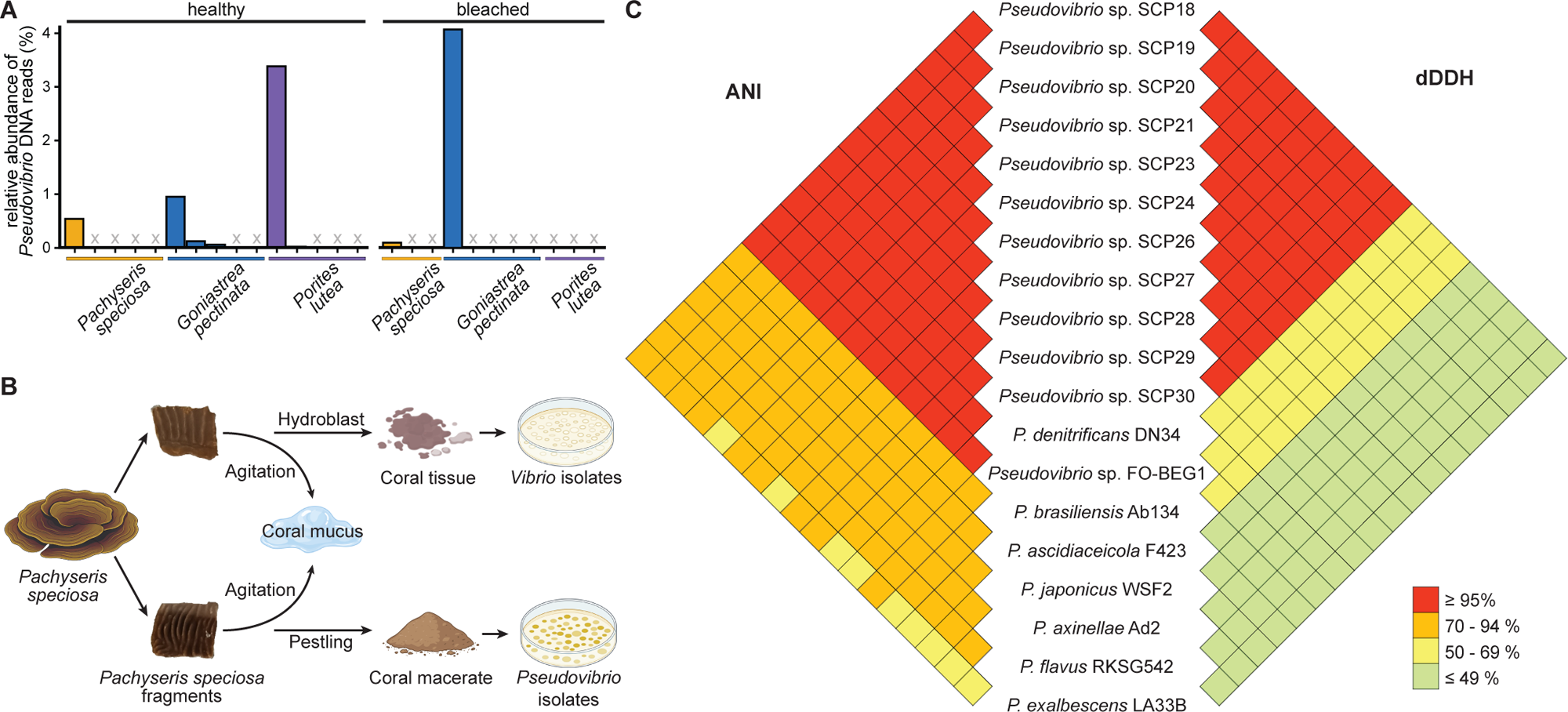
Singaporean corals show scattered distribution of *Pseudovibrio* in healthy and bleached states, and *Pseudovibrio* isolates from *P. speciosa* have almost identical genomes. (A) Relative abundance (%) of *Pseudovibrio* reads in *P. speciosa*, *G. pectinata*, and *P. lutea* in healthy and bleached corals demonstrate changes between coral species and coral health conditions. (B) Isolation of pure cultures of *Vibrio* and *Pseudovibrio* isolates from *P. speciosa* tissue and macerate respectively. (C) Pairwise comparisons of *Pseudovibrio* genomes by average nucleotide identity (ANI) and digital DNA-DNA hybridisation (dDDH) scores. Genome-sequenced *Pseudovibrio* type strains, and the isolate FO-BEG1 from a black-band diseased scleractinian coral, were used for comparisons.

### Near-clonal Pseudovibrio isolates differ by carriage of the chromid pCJH

Eleven *Pseudovibrio* isolates with brown colonies were obtained from coral macerates of *P. speciosa* (Fig. 1B). All isolates shared > 99.99% ANI (sequence identity of shared genomic regions) with each other (Fig. 1C), indicating they belong to a single species. Comparison to the closest related genome (MiGA), *Pseudovibrio* sp. FO-BEG1 (Fig. S2), and to all *Pseudovibrio* type strains, the closest being *P. denitrificans* DN34, gave ANI > 95% but dDDH < 70% (Fig. 1C). Therefore, isolates were identified to genus, not species level (subsequently referred to as *Pseudovibrio* sp. SCP18-21, 23-24, 26-30).

Genome comparison showed that *Pseudovibrio* sp. SCP18, SCP21, SCP24, SCP27, SCP28, SCP29 and SCP30 have an additional ∼490 kbp contig that is absent in SCP19, SCP20, SCP23 and SCP26 (Fig. 2A, B). These four isolates without the contig have a 43 bp region with 95.3% identity to an intergenic spacer with inverted repeats on the large contig. No plasmids were detected with PlasmidFinder, but manual inspection for plasmid origin of replication sites identified a *repABC* operon in the contig. Given its large size, GC content (52.7%) similar to that of the chromosome (52.2%), strongly correlated relative synonymous codon usage (RSCU) with the chromosome (Pearson’s r = 0.98), together with the presence of three essential genes (*map*, *fabI*, *dnaK*; Fig. 2B), this contig represents a putative chromid [51]. This is subsequently referred to as pCJH, reflecting the climate-driven Jekyll-and-Hyde phenotype of opportunistic pathogen and beneficial symbiont mediated by the chromid.

**Figure 2:**
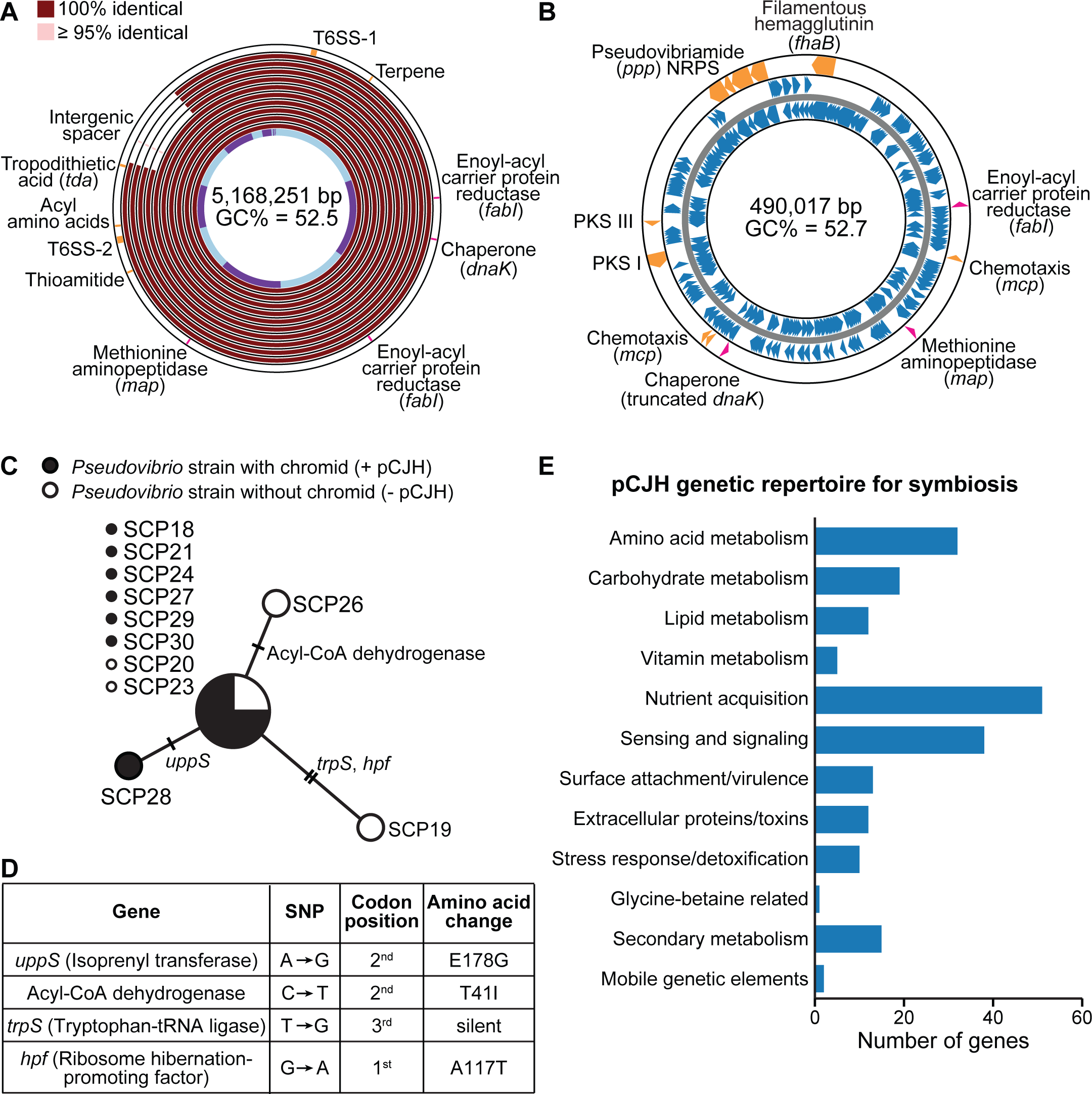
The chromid pCJH distinguishes *Pseudovibrio* strains and carries putative symbiosis-relevant genes. (A) BLAST atlas of 11 genomes identified a ∼490 kbp contig present in 7 of the genomes. Genome contigs are shown in light blue and purple on the inner track. Core essential genes (*map*, *fabI*, *dnaK*) are labelled in pink and genes of interest in orange on the outer track. Colours on each track indicate BLAST hits on 10 genomes and identity relative to the SCP18 reference genome. The four isolates without chromid have a 43 bp region with 95.3% similarity to an intergenic spacer on the SCP18 additional contig. (B) Manual examination identified this contig as a *repABC* chromid (pCJH), with similar GC content and codon usage bias to the chromosome. Secondary metabolite gene clusters and mechanisms that are suggested to be involved in symbiosis are shown in orange, as well as the essential core genes *map*, *fabI*, and *dnaK* in pink. Coding sequences are shown in blue. (C) Minimum spanning tree of the 11 *Pseudovibrio* genomes based on the 4 identified single nucleotide polymorphisms (SNPs). Node size reflects the number of isolates sharing an identical genotype, and branch labels indicate the gene containing each SNP, with tick marks denoting the number of nucleotide differences. (D) SNPs identified in each gene, with the corresponding codon position and resulting amino acid change. (E) The genes carried on pCJH were assigned to symbiosis-relevant categories by manual curation, cross-referencing annotations from eggNOG-mapper, Prokka, and BLASTp.

Four single nucleotide polymorphisms (SNPs) were identified among the 11 genomes (Fig. 2C), both in strains with and without the chromid, with no fixation of the chromid in the populations. The SNPs were localised to the *trpS, hpf, uppS* genes and an uncharacterized acyl-CoA dehydrogenase (Fig. 2D), none of which fall within characterised active or catalytic site. No SNPs were identified among pCJH sequences.

Gene clusters found by antiSMASH in the chromosome include thioamitide, acyl amino acids and terpene biosynthesis (Fig. 2A). The *tda* gene cluster and two type six secretion systems (T6SSs) were identified through BLASTp on the chromosome. Analysis of the SCP18 pCJH revealed functions that could support *Pseudovibrio* symbiosis and persistence within the coral host-associated environment (Fig. 2E and S3; Supplementary Note 1). The chromid carries genes associated with amino acid, carbohydrate, lipid and cofactor/vitamin metabolism, together with 51 predicted nutrient and metabolite transport proteins, including 5 complete TRAP-type transporters. It also harbours two polyketide synthase gene clusters (PKS I and PKS III), and a large NRPS gene cluster (Fig. 2B) homologous to the experimentally characterized *P. brasiliensis ppp* locus, whose products, pseudovibriamide A and B, modulate motility, biofilm formation and overall gene expression [52].

### Chromid pCJH affects Pseudovibrio growth rate, cell density, and attachment to corals

To assess if the chromid expanded *Pseudovibrios* genetic repertoire for its interaction with *P. speciosa* and its physiology, SCP18 (with chromid, pCJH) and SCP20 (without chromid) were chosen as representative strains as they have identical genomes (i.e. no SNPs).

Comparison of growth curves between *Pseudovibrio* strains (Fig. S4) showed significant differences between isolates with and without the chromid (Mann-Whitney rank sum test, p < 0.001) (Fig. 3D), with a specific growth rate of 0.30 h^-1^ without the chromid and 0.17 h^-1^ with the chromid. Additionally, they reached significantly different stationary phase cell density (t-test, p = 0.01) and CFU of 1.23 × 10^10^ cells mL^-1^ at 12 h and 3.02 × 10^9^ cells mL^-1^ at 16 h without and with the chromid respectively.

**Figure 3:**
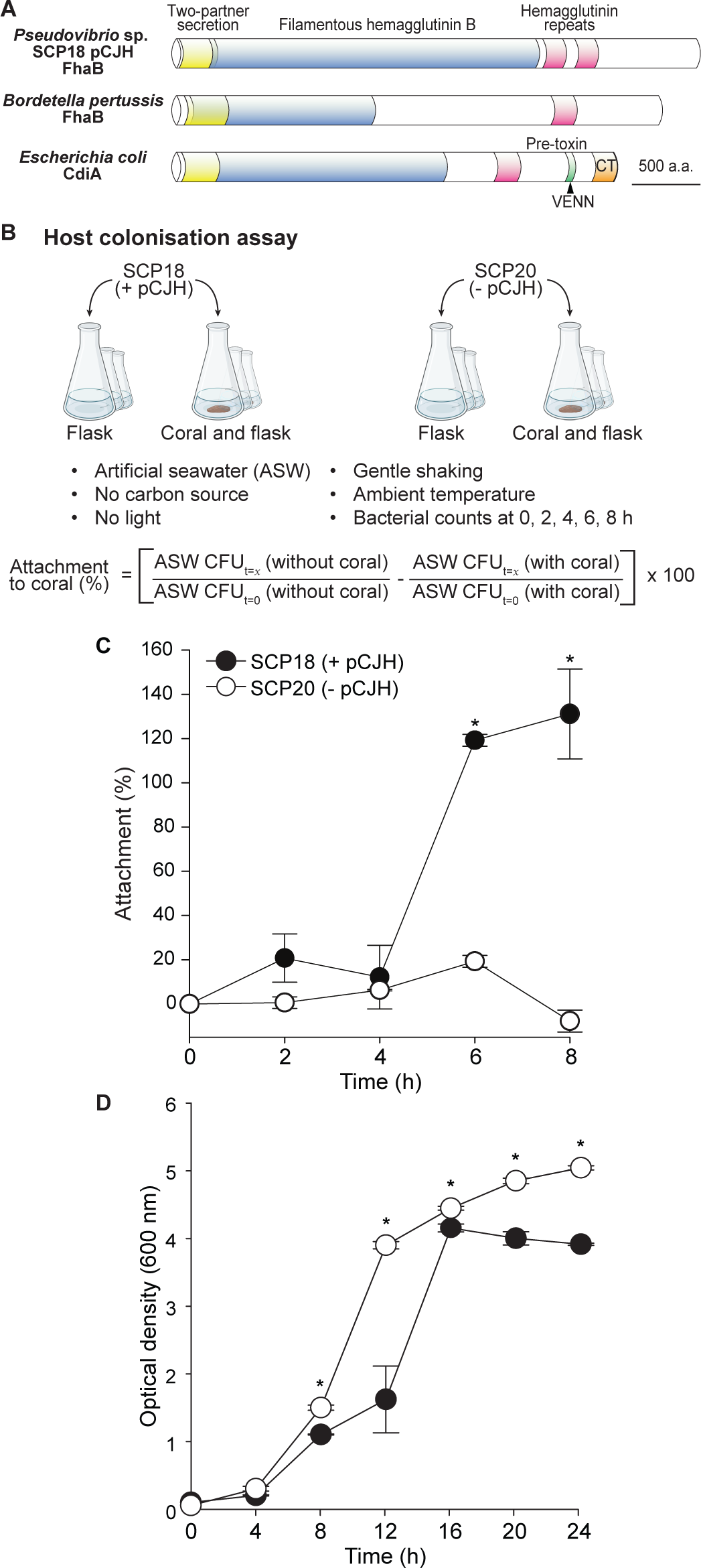
Chromid pCJH promotes attachment to coral host, and imposes a cost on bacterial growth. (A) Domain architecture of FhaB, a filamentous adhesin compared to its homolog in *Bordetella pertussis* and to CdiA, the two-partner secretion protein of *Escherichia coli* contact-dependent growth inhibition. FhaB lacks the VENN motif and C-terminal toxin domain that enable CdiA-mediated killing. (B) Schematic of the host colonisation assay, in which attachment of *Pseudovibrio* isolates SCP18 (with pCJH) and SCP20 (without pCJH) to *P. speciosa* was measured over 8 h. Independent replicates (i.e. three flasks with three separate coral fragments, and three flasks without coral fragments) were set up for each bacterium. The total number of samples was: n = 4 (conditions) × 3 (replicates) × 5 (timepoints) = 60. (C) Attachment to *P. speciosa* by SCP18 and SCP20 over 8 h. (D) Optical density (OD_600_) of SCP18 and SCP20 cultures over 24 h. Error bars indicate standard error. Statistical comparisons were made using Mann-Whitney rank sum test.

Although the core motility and chemotaxis machinery is chromosomally encoded, pCJH expands the sensory repertoire with 3 additional methyl-accepting chemotaxis proteins (MCPs) and a PAS-domain sensory protein potentially related to an aerotaxis receptor, complementing the 15 chromosomal MCPs. It also encodes a multidomain c-di-GMP signalling protein, potentially integrating motility, chemotaxis and surface attachment.

Notably, pCJH encodes a CdiA-like protein with high amino acid identity to filamentous hemagglutinin (FhaB) protein lacking a toxin domain (Fig. 3A), characteristic of an adhesin for surface attachment [53]. Also on pCJH are components of capsular polysaccharide biosynthesis and export, analogous to those in rhizobia required for host attachment [54].

To test whether these chromid-encoded functions translate into greater host association, we compared the attachment of SCP18 (with pCJH) and SCP20 (without pCJH) to *P. speciosa* fragments in a host colonisation assay (Fig. 3B). At 6 and 8 h, attachment to *P. speciosa* was significantly higher for SCP18 than for SCP20 (Mann-Whitney rank sum test, p < 0.001), with SCP18 increasing from 12 ± 14% to 119 ± 3% at 6 h (Fig. 3C), implicating pCJH in enhanced host colonisation. Attachment above 100% likely reflects growth on organic carbon released by respiration and leakage, processes that the dark assay does not limit, unlike photosynthesis.

### Increase in coral thermal resilience from pCJH

Inoculation with SCP18 or SCP20 altered host outcomes relative to uninoculated *P. speciosa* under temperature ramping but not under constant temperature (Fig. 4A). Constant temperature elicited no observable or statistical effects on any treatment group (Fig. 4B).

**Figure 4:**
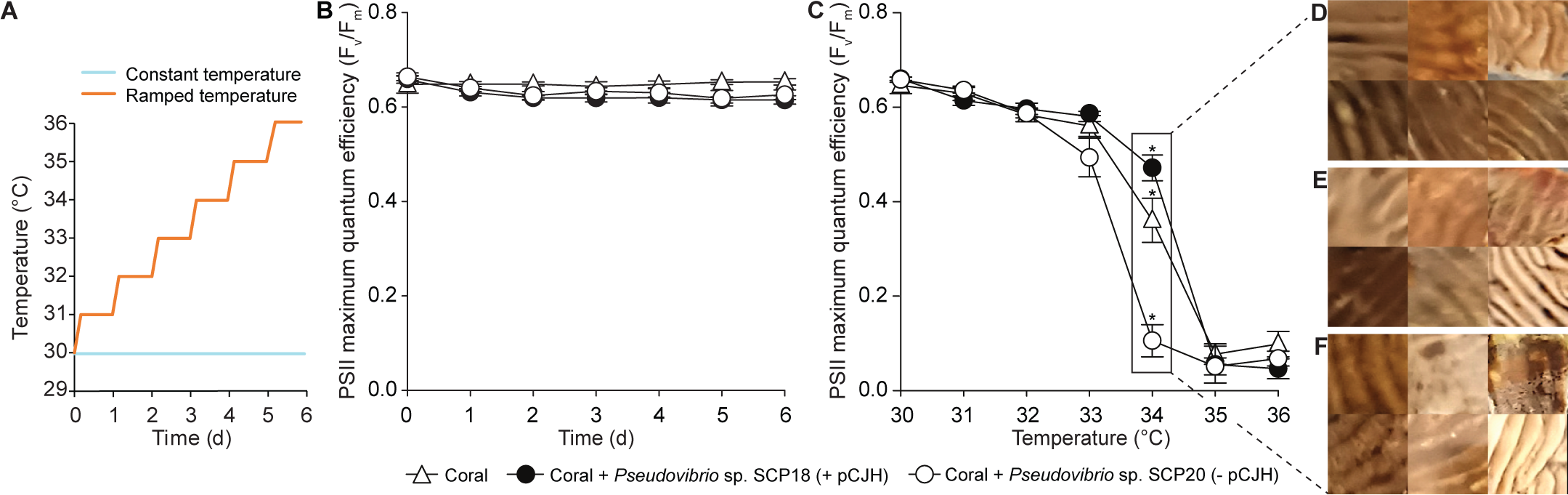
*Pseudovibrio* isolates without the pCJH chromid reduces thermal tolerance, and isolates bearing pCJH increase it. (A) *P. speciosa* fragments inoculated with SCP18 (with pCJH) and SCP20 (without pCJH), as well as control fragments without any inoculated bacteria, were exposed to ramped-temperature treatment (1 °C increase per day) or maintained at constant temperature (30 °C). PAM fluorometry was used to measure photosystem health (photosystem II maximum quantum efficiency, F_v_/F_m_). (B) Coral host F_v_/F_m_ under constant temperature regime. (C) Coral host F_v_/F_m_ under ramped-temperature treatment. Error bars indicate standard error. Statistical comparisons were made using ANOVA with Holm-Sidak test (p ≤ 0.05). Samples used in this aquaria experiment: n = 3 treatments (no inoculation, SCP18, SCP20) x 2 temperature regimes (constant and ramping) x 6 replicate coral fragments = 36. Images of coral fragments with (D) SCP18 (with pCJH), (E) no treatment, and (F) SCP20 (without pCJH) corroborate these findings.

Under ramping, uninoculated fragments reached a tipping point, defined as the last temperature before F_v_/F_m_ dropped significantly at the following temperature, at 33 °C (Fig. S5). Fragments inoculated with SCP20, lacking pCJH, reached their tipping point also at 33°C, whereas fragments inoculated with SCP18, carrying pCJH, reached theirs at 34 °C (Fig. 4C, S5). At 34 °C, F_v_/F_m_ was significantly lower in corals inoculated with SCP20 (Holm-Sidak, p < 0.001) and significantly higher in SCP18-inoculated corals (Holm-Sidak, p = 0.05) than in uninoculated corals (Fig. 4C). Bleaching, most evident in corals treated with SCP20 and least in those treated with SCP18, was first observed on ridges and progressed into depressions (Fig. 4D–F).

### Changes in the coral microbial community in response to temperature ramping

Temperature ramping experiments performed on four coral species without the addition of *Pseudovibrio* isolates were carried out to investigate bacterial community changes using 16S metabarcoding. The relative abundances of the second most abundant ASV, ASV0002 (100% identity with *Vibrio* sp. SCP2 and *V. coralliilyticus* SCP4), and ASV0135 (100% identity with *Pseudovibrio* sp. SCP18-21, 23-24, 26-30) were not significantly impacted by the heat treatment (Fig. 5A). Only ASV0006 (100% identity with *V. harveyi* SCP5) had a significant (t-test, p = 0.01) decrease in proportional abundance under the heat treatment.

**Figure 5:**
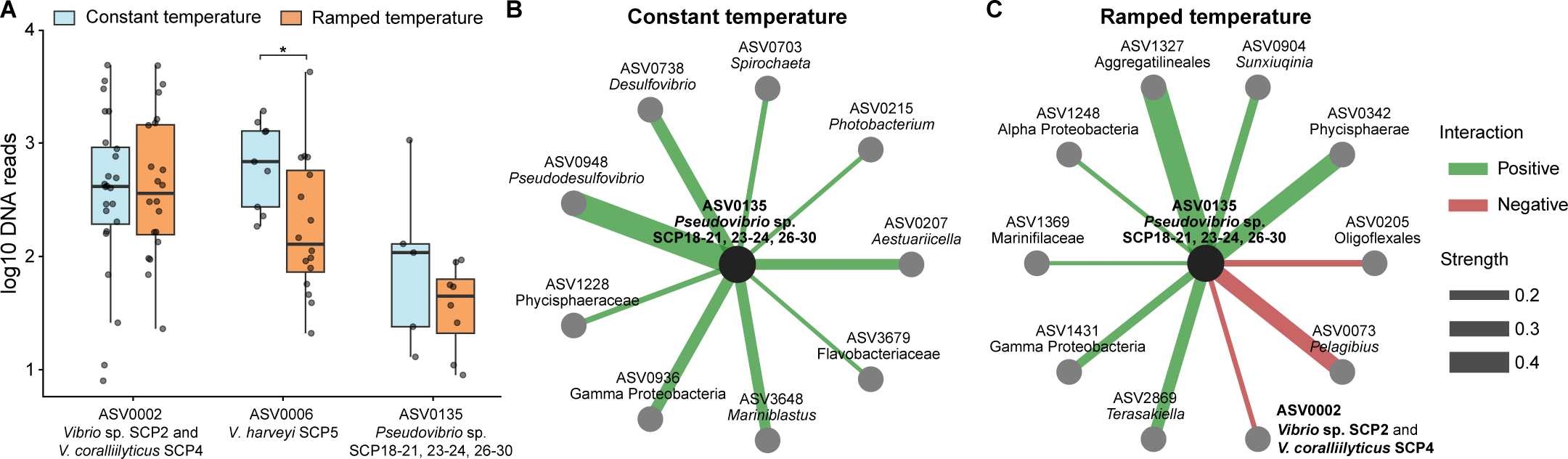
*Pseudovibrio* ASVs corresponding to SCP isolates are present in multiple corals species, even after ramped temperature treatment, and display negative interactions to local *Vibrio* strains. Total samples used in this experiment n = 4 coral species x 2 heat treatments x 6 coral fragments = 48. The (A) relative abundance of ASVs that have 100% sequence identity with SCP strains, under constant and ramped temperature treatments. Boxplots centre line indicates the median, with box limits encompassing between and whiskers extending to 1.5x interquartile range. Asterisk denotes statistically significant difference using Student’s t-test (p < 0.05). SPIEC-EASI network models for coral fragments under (B) constant and (C) ramped temperature treatments showing positive (green) and negative (red) interactions between ASV0135 (*Pseudovibrio* SCP strains) and other ASVs. Edge thickness indicates interaction strength.

The *Vibrio* ASVs were the second and sixth most proportionally abundant ASVs in this experiment. ASV0135 had the greatest proportional abundance among *Pseudovibrio* ASVs and was patchily distributed, present in 5 of 24 control and 8 of 21 heat-treated fragments (Fig. S6).

Network analysis of ASVs from samples in the control set up identified only positive associations with ASV0135 *Pseudovibrio* sp. (Fig. 5B). Interestingly, under heat ramping conditions producing bleaching disease, negative interactions were detected, including one with the highly abundant ASV0002 *Vibrio* (Fig. 5C). Positive interactions were also identified, however, none were common to both bacterial communities under constant and ramped temperature regimes.

### Contact-dependent and -independent inhibition of Vibrio species by Pseudovibrio isolates

Because *Vibrio* spp. are implicated in bleaching disease, we tested whether the *Pseudovibrio* isolates inhibit *Vibrio* isolated from the same host. All *Pseudovibrio* isolates inhibited *Vibrio* isolates (Fig. 6A), producing measurable zones of inhibition (ZOIs) with no variation between replicates of the bioassay (Table S3). A notable pattern emerged where activity against the two *V. coralliilyticus* isolates and the unclassified *Vibrio* sp. SCP2 showed relatively small ZOIs with a dense lawn at the edge of the ZOI (Fig. 6B), compared to the larger ZOIs against *V. harveyi* strains with a diffuse edge to the ZOI (Fig. 6C). However, there was no apparent separation in activity between strains with or without pCJH, nor between strains with and without SNPs (Fig. 6D). AntiSMASH and manual searches identified many candidate diffusible antibiotics in both the chromosome and pCJH (Fig. 6E). On the chromosome, searches for contact-dependent mechanisms identified two T6SSs present in all genomes, which are the likely mechanism, although an unknown contact-dependent mechanism cannot be excluded.

**Figure 6:**
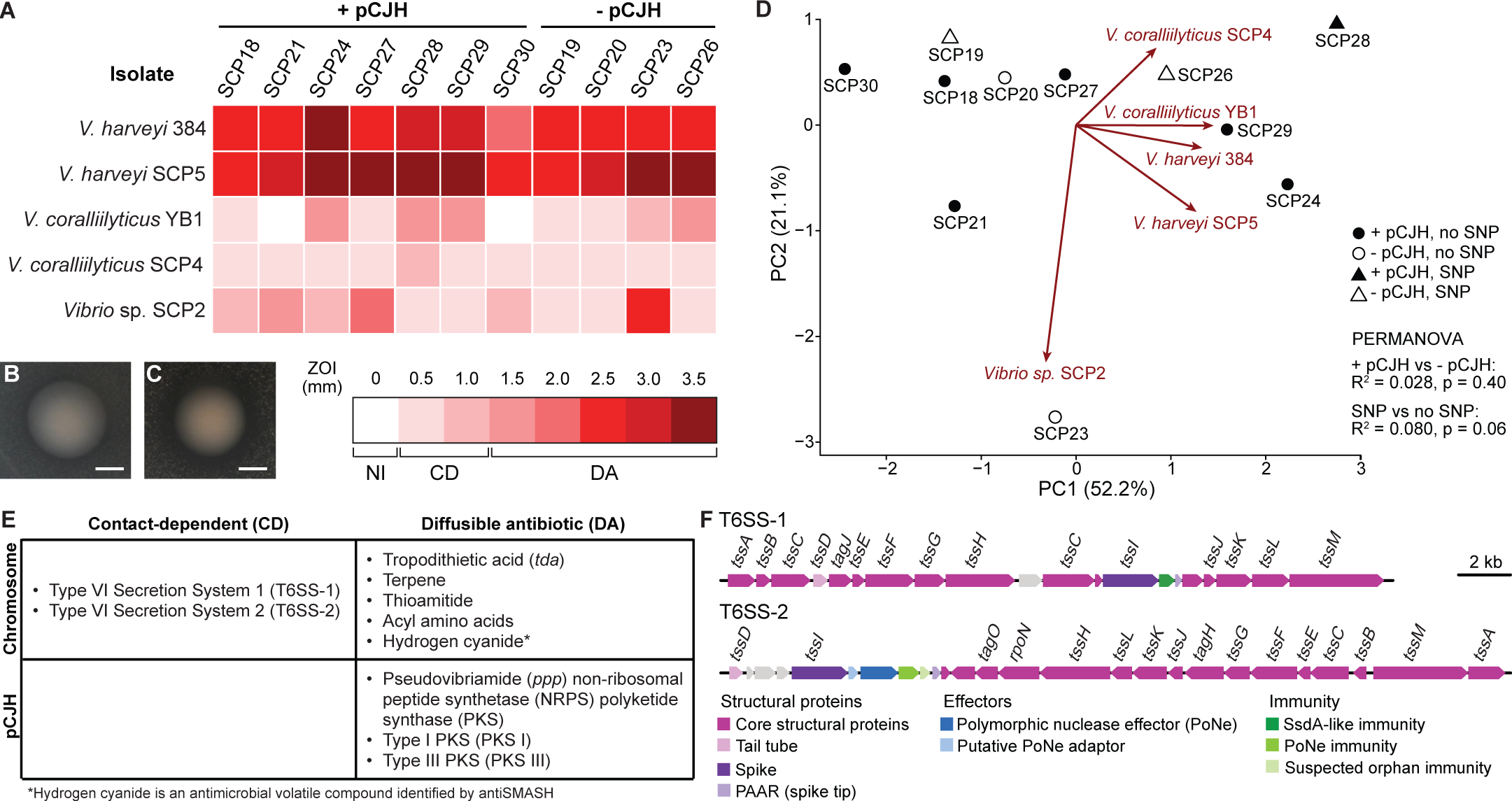
*Pseudovibrio* isolates with and without the pCJH chromid equally inhibit *Vibrio* isolates, with T6SSs implicated in contact-dependent inhibition. (A) *Pseudovibrio* isolates were spotted on lawns of five *Vibrio spp.* isolates, and zones of inhibition (ZOI) were measured and shown as a heat map. Aside from no inhibition (NI), two types of inhibition were observed: (B) contact-dependent (CD) inhibition, and (C) inhibition from a diffusible antibiotic (DA). (D) A principal component analysis of the ZOI showed no significant differences (PERMANOVA, p > 0.05) between strains with and without the pCJH, or with and without SNPs. (E) antiSMASH and manual searches of both the chromosome and chromid identified T6SSs as primary candidates for contact-dependent inhibition, and a repertoire of diffusible antibiotics, encoded on both the chromosome and pCJH, as primary candidates for diffusible-antibiotic inhibition. (F) Gene clusters encoding the two T6SSs, T6SS-1 and T6SS-2, found across all *Pseudovibrio* isolate genomes, with genes colored by category.

### T6SS as a potential contact dependent killing mechanism for Pseudovibrio

Two divergent T6SS loci (T6SS-1 and T6SS-2; Fig. 6F), potentially enabling contact-dependent inhibition, are located in all genomes. The gene order of both loci is non-syntenic and not completely overlapping, yet together they encode all genes involved in building the canonical T6SS apparatus (*tssA*-*tssM* and *vgrG*, coding for the trimeric T6SS spike complex) [55]. T6SS-1 is highly conserved among *Pseudovibrios* and encodes a putative DUF6931/SsdA-like immunity protein [56] and a PAAR (i.e. spike tip) protein downstream of *vgrG*, while notably lacking the typical *hcp* (tail tube) gene [55]. T6SS-2 encodes the missing *hcp* gene but lacks the *tssA* gene coding for the tube cap protein present in T6SS-1. It differs from otherwise similar *Pseudovibrio* T6SS loci in the variable effector-immunity gene region downstream of *vgrG*, which encodes a polymorphic nuclease effector protein [57] flanked by putative cognate adapter and immunity proteins, followed by a suspected orphan immunity protein (Supplementary Note 2). The variable region ends with a hypothetical + PAAR-protein pair also found in many other *Pseudovibrios* with other effector-immunity gene arrays.

## Discussion

We have shown that coral-associated *Pseudovibrio* from equatorial turbid reefs can alter *P. speciosa*’s resilience to temperature stimulated bleaching. Its chromid, pCJH, genetically underpins a novel, climatically impacted Jekyll-and-Hyde interaction, in which *Pseudovibrio* is an opportunistic pathogen without pCJH and a symbiont with it, extending the coral’s thermal resilience to bleaching (Fig. 7). This transition of ecological role contributes to coral resilience in Singapore’s extreme reefs.

**Figure 7:**
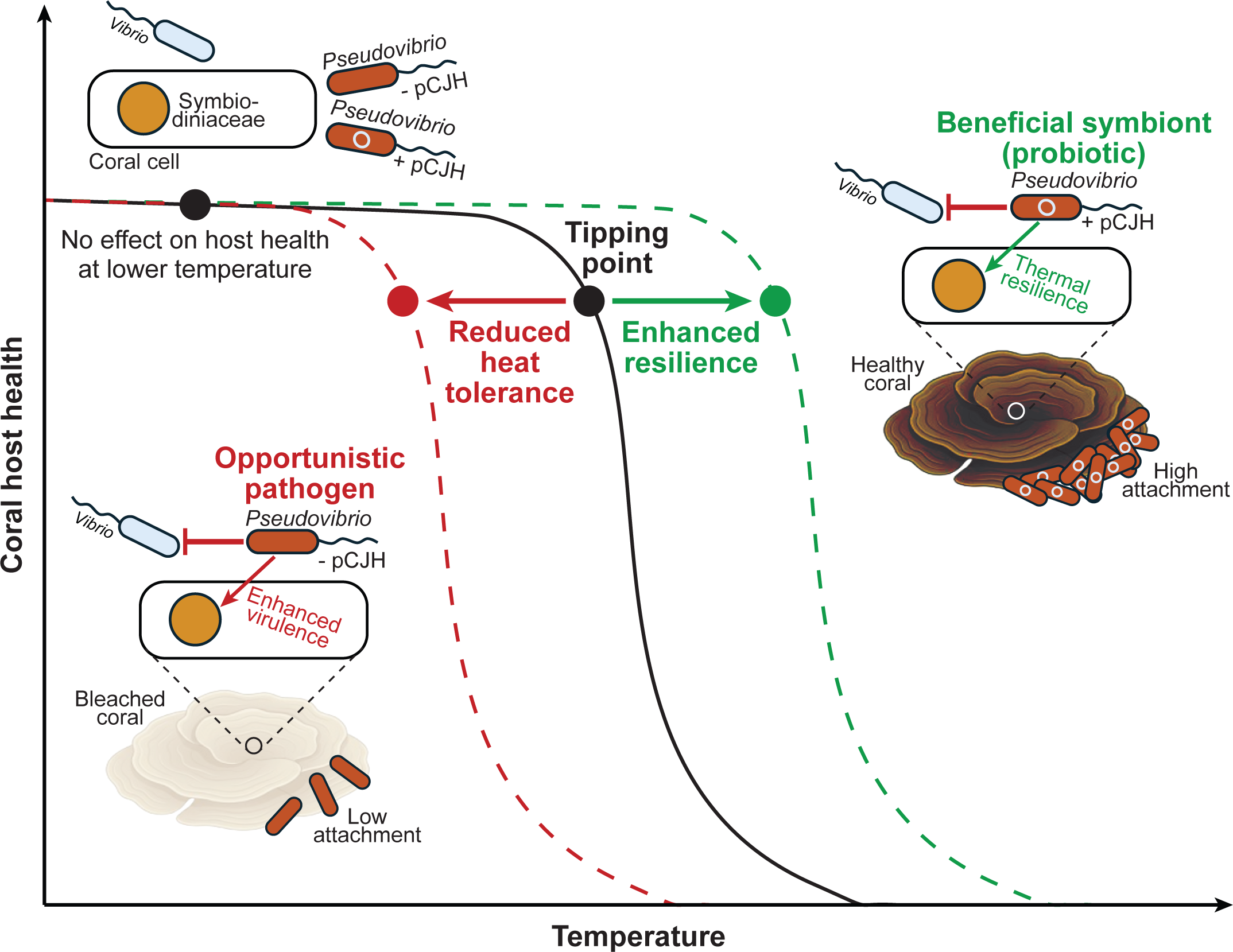
The chromid pCJH determines whether a novel *Pseudovibrio* species acts as a coral symbiont or pathogen under heat stress. At lower temperatures, coral host health is unaffected by pCJH carriage. *Pseudovibrio* isolates inhibit *Vibrio* pathogens regardless of pCJH carriage. *Pseudovibrio* isolates carrying pCJH act as beneficial symbionts, conferring enhanced attachment to the coral host and extending the thermal range over which the coral’s Symbiodiniaceae maintain photosynthetic health. Conversely, isolates lacking pCJH act as opportunistic pathogens by exacerbating PSII decline beyond this tipping point.

Members of the genus *Pseudovibrio*, widely distributed marine bacteria frequently associated with sponges, corals and other invertebrates [58, 59], are candidate beneficial symbionts to their hosts due to their genomic potential for host interaction and symbiosis (including the machinery for producing bioactive compounds), being correlated with healthy coral hosts [60–62]. Consistent with this broad host range, we detected *Pseudovibrio* in all four coral species examined in the field and acute heat experiments, although its relative abundance and prevalence differed among hosts (Fig. 1A, 5A and S6). Host-specific response to bleaching suggests that *Pseudovibrio* is a flexible rather than fixed member of the coral microbiome, and that its relationship with the host shifts with environmental conditions, a property relevant to its potential use as a BMC.

However, our isolates show that these ecological roles can vary widely even within a single *Pseudovibrio* clonal complex: genomic divergence among the 11 isolates is extremely low (four SNPs in total; Fig. 2C), with strains segregated only by the presence or absence of a ∼490 kbp chromid present in 7 of 11 genomes (Fig. 2A). Notably, strains carrying pCJH are phenotypically distinct, exhibiting enhanced colonisation of *P. speciosa* (Fig. 3C) and altered host interactions (Fig. 4) compared with chromid-free strains. The four SNPs localised to *trpS*, *hpf*, *uppS* and an uncharacterized acyl-CoA dehydrogenase (Fig. 2C) fall in genes unrelated to adhesion, secretion or secondary metabolism (Fig. 2D) and are unlikely to explain these differences, implicating pCJH as the primary determinant of these phenotypes.

Megaplasmids and chromids encode and mobilise accessory functions (e.g. adhesins, secretion and regulatory modules, biosynthetic gene clusters) that can drive large phenotypic shifts without chromosomal divergence [63, 64]. Variable accessory replicons are reported across *Pseudovibrio* genomes, and our nearly clonal isolates with and without pCJH show that chromid carriage can be dynamic at the population level rather than fixed.

Interestingly, *Pseudovibrio* sp. FO-BEG1 carries a 441 kb chromid with 94.8% ANI with pCJH. The fact that the *Pseudovibrio* SCP isolates and FO-BEG1 were both isolated from corals and harbour chromids suggests that the chromid is stable across geographical and temporal scales and may be ecologically important for association with corals. Persistence of these costly extrachromosomal elements should reflect context-dependent benefits that outweigh maintenance costs [65]. We demonstrated that maintaining the chromid imposes a physiological cost, reducing both growth rate and population density (Fig. 3D), the classical fecundity cost that drives genome streamlining [65]. In addition, analysis of sequence coverage indicates a 1:1 chromosome:chromid copy-number, consistent with chromosomal-style replication and supporting the high energetic cost of maintaining pCJH.

Despite these costs, we experimentally demonstrated that pCJH provides at least two selective advantages: enhanced host colonisation through increased attachment to coral fragments (Fig. 3C), and improved host photophysiology and pigmentation under heat stress (Fig. 4C, D). These benefits are consistent with pCJH encoding functions commonly found on marine symbiont extra-chromosomal elements (adhesins/TPR/TPS motifs, polysaccharide biosynthesis and export, chemotaxis receptors and PKS/NRPS biosynthetic gene clusters) that promote colonisation and bioactive compound production [60, 66]. Genes that could help *Pseudovibrio* mitigate reactive chemical stress [67, 68] in the coral holobiont are also found on the chromid. These include genes for formaldehyde, methylglyoxal (by-product of Calvin cycle in stressed photosymbionts) and nitric oxide detoxification, peroxidases and a glutathione-dependent detoxification system, consistent with the proposed role of bacterial ROS-scavenging functions in mitigating oxidative stress within coral holobionts [67–70]. Although candidate genes for glycine betaine uptake, and downstream sarcosine metabolism are on the chromosome, the chromid has a key gene, dimethylglycine dehydrogenase, linking pCJH to glycine-betaine turnover, important for osmoprotection and nitrogen cycling [71].

Carbohydrate-specific binding of *V. coralliilyticus* enables infection from just 100 cells mL^-1^ [49]. Meanwhile, enhanced colonisation will increase proximity-dependent interactions in the host tissue microenvironment, such as metabolite exchange and contact-mediated inhibition which can restructure the holobiont’s microbial community [49, 72, 73]. Drawing parallels to our experimental results, pCJH-enhanced colonisation of *Pseudovibrio* likely underlies its improvement of coral photophysiology under heat stress (Fig. 4C, D).

Conversely, *Pseudovibrio* without pCJH significantly reduced Symbiodiniaceae PSII function under ramped temperature, causing a greater decline in PSII function and more severe bleaching beyond the tipping point (Fig. 4C, F), but had no detectable effect at constant temperature (Fig. 4B). We therefore describe it as an opportunistic pathogen in the sense of the damage-response framework, in which pathogenicity is defined by host damage and depends on host state rather than being an intrinsic property of the microbe [74].

Because SCP18 and SCP20 share no chromosomal SNPs (Fig. 2C), our experiments provide evidence that pCJH-encoded functions are the proximate cause of transitioning *Pseudovibrio* from an opportunistic pathogen to a beneficial coral symbiont under heat stress conditions. This genetic fluidity, and the phenotypic plasticity it confers, is critical to survivorship under environmental change. Under a ramped temperature regime that destabilises the photosymbiont-coral symbiosis and reshapes bacterial interactions [75], this plasticity, embodied by a large mobile genetic element such as pCJH, is a survival strategy that can extend thermal resilience of a coral and its holobiont.

T6SSs are contact-dependent nanomachines that inject toxic effector proteins into neighbouring cells, shaping niche colonisation and competitive fitness, consequently altering community structure in marine systems [76–78]. They are especially effective in biofilms or host mucus where cells are closely packed, and elevated temperatures can increase T6SS-mediated bacterial virulence [79]. All isolates encode two chromosomal T6SS loci (T6SS-1 and T6SS-2; Fig. 6F), and the observed contact-dependent inhibition across both pCJH-bearing and pCJH-free strains (Fig. 6A) implicates these chromosomal systems as likely mediators of proximity-dependent antagonism. By contrast, pCJH-encoded factors like pseudovibriamides offer *Pseudovibrio* a combinatorial approach to the chemical arms race (i.e. diffusible antagonism) in that it promotes swimming, a planktonic lifestyle, reducing both colonisation and biofilm-formation and thereby could enhance antibiotic effectiveness [80].

Network analysis revealed a reshape of bacterial interactions under coral heat stress. The *Pseudovibrio* ASV0135 (including all isolates with and without pCJH for *Pseudovibrio*) exhibited only positive associations within the coral microbiome under control conditions (Fig. 5B), consistent with a cooperative bacterial community. In contrast, heat stress introduced negative associations, notably between *Pseudovibrio* and *Vibrio* ASVs, and its positive associations shifted to different ASVs (Fig. 5C). ASV0006 (*V. harveyi* SCP5) shows a significant decrease in proportional abundance under heat stress (Fig. 5A), contrary to the usual heat-driven increase in *Vibrio* relative abundance [81, 82], indicating the emergence of possibly antagonistic interactions. These known coral pathogens upregulate virulence at elevated temperatures [24, 49], yet our study adds community context by showing how heat reshapes bacterial interactions concurrently. Mechanistically, these shifts can reflect increased activity of contact-dependent and diffusible antagonism mechanisms [77, 78, 83]. Thus, holobiont outcomes under heat stress emerge from both direct pathogen virulence against the coral photosymbiont and from restructured bacteria-bacteria antagonism, which is a dual pathway that could explain how microbiome composition and interaction networks predict coral thermal responses [17, 75].

The ecological plasticity conferred to this previously undescribed *Pseudovibrio* species by pCJH underscores the dynamic nature of these interactions. In *Phaeobacter inhibens*, algal host cues trigger a switch from symbiosis to pathogenic interaction [84, 85]. Our results demonstrate a novel, Jekyll-and-Hyde switch in the reverse direction, in which pCJH shifts this *Pseudovibrio* species from an opportunistic pathogen to a symbiotic partner in a tripartite relationship with the photosymbiont and coral. Interaction analysis suggests this symbiosis is further stabilized by increasing bacterial-bacterial interaction complexity as the primary photosymbiont-coral symbiosis becomes stressed under elevated temperature. Acquisition of extrachromosomal elements is known to enable commensal bacteria to become symbionts, as in rhizobia, where symbiotic megaplasmid (pSym) carry genes essential for host-specific associations [86, 87]. The *Pseudovibrio* chromid pCJH represents a conceptually similar, but novel, genetic determinant of host association. This points to a broader role for mobile genetic elements in coral reef resilience, both in identifying other candidate BMCs with similar genomic architecture, and in using beneficial mobile genetic elements for corals (BMGECs) to enhance coral resilience.

## Acknowledgements

We thank Huang Danwei and Yann Boucher from the National University of Singapore for their valuable feedback on the manuscript. The authors would like to acknowledge St. John’s Island National Marine Laboratory (SJINML) for providing the facility necessary for conducting the research. The Laboratory is a National Research Infrastructure under the National Research Foundation Singapore. We would also like to thank the SJI Coral team at the NUS Tropical Marine Science Institute (TMSI) for collecting the coral fragments used in this research (NParks permit no.NP/RP22-127-2). Any opinions, findings and conclusions or recommendations expressed in this material are those of the author(s) and do not reflect the views of National Research Foundation, Singapore and National Parks Board, Singapore.

## Author contributions

LKD, JTIT and RJC conceived the project; RJC, JPAP, JCHK, WL, JCHK, LCSN & JTIT contributed to experimental design; JPAP, JJHL, PM, WL, JCHK, IT and LJWL performed the experiments; JPAP, AARL, JJHL, PM, IT and LJWL generated the data; CWHS, HHCL, JPAP, SS, AARL, JJHL, PCK and RJC analysed the data; CWHS, HHCL, JPAP, SS, PCK and RJC contributed to figure preparation; CWHS, HHCL, JPAP, SS, PCK, LCSN, JTIT and RJC contributed to writing the manuscript.

## Data availability

The Sequence Read Archive (SRA) data was deposited to NCBI, and the complete genome sequences was deposited to GenBank under accession numbers listed in Table S1.

