## Supplementary Material for "Chromid transforms opportunistic pathogen into beneficial symbiont to raise coral’s thermal tipping point"

**Supplementary materials and methods**

Molecular community and interaction analyses

Extracted DNA was stored at -20 °C. DNA concentration was quantified using a Qubit™ 2.0 Fluorometer in conjunction with the Qubit™ dsDNA Broad-Range Assay Kit (Thermo Fisher Scientific, Waltham, USA). Samples that yielded DNA concentrations lower than 1 ng/µL (i.e. *P. acuta*) were excluded from further processing as they did not provide the minimum amount (20 ng) of template DNA for PCR.

PCR was performed in 25 µL containing 10 µM of each primer, 1× KAPA Plant PCR Buffer, 1 U of KAPA3G Plant DNA polymerase (Roche, Basel, Switzerland), 2.25 µg of bovine serum albumin (BSA), and 20-50 ng of template DNA. PCR products were visualised on a 1% agarose gel. Bands corresponding to ~360 bp were excised and purified using the Wizard® SV Gel and PCR Clean-up System (Promega, Madison, USA), and quantified with a Qubit™ 2.0 Fluorometer using the Qubit™ dsDNA High-Sensitivity Assay Kit (Thermo Fisher Scientific, Waltham, USA).

All bioinformatic and statistical analysis for 16S metabarcoding were performed in R v4.4.2 [1]. *Cutadapt* v4.0 [2] was used to remove primers and adapters from demultiplexed sequences. Raw sequence quality was assessed using the *plotQualityProfile* function from the *DADA2* package v1.34.0. [3] and truncation lengths were selected based on where the median quality score remained above the minimum threshold of Q30. Forward and reverse reads were trimmed between 240 and 160 base pairs respectively with the *filterAndTrim* function, and a maximum expected error of 2 was used for both forward and reverse reads. Default values were used for all other parameters. Error rates were estimated *de novo* from the filtered reads using the *learnErrors* function, which were then used to infer true sequence variants. Forward and reverse reads were subsequently merged using the *mergePairs* function to form amplicon sequence variants (ASVs), and chimeric sequences were removed using the *removeBimeraDenovo* function with the consensus method. Upon taxonomic assignment, Chloroplast and mitochondria sequences were removed, and samples normalised to the median read count (30,985 reads) of the filtered dataset.

BSL2 Aquaria experimental setups

The Biosafety Level 2 aquaria at SJINML was used in the experiment “Pseudovibrios interaction with temperature stressed coral” as detailed in the main manuscript’s Methods section. The aquaria utilised sand-filtered seawater cycled through a 100 µm filter, using Evergrow IT AquaOcean IT5080 lights on a 12:12 h (light:dark) diurnal cycle at 120 µmol $m^{-2}$

$s^{-1}$ photosynthetically active radiation (PAR), measured using an Apogee MQ-510 underwater quantum meter to ensure consistent illumination across tanks. Corals were imaged daily, during the first hour of the light period, when temperature readings were taken using a YSI ProQuatro probe.

BSL1 Aquaria setup

The Biosafety Level 1 aquaria at SJINML was used in the experiment "*Heat stressed corals' impact on bacterial-bacterial interactions*" as detailed in the main manuscript's Methods section. *T*welve fragments of each coral species were tested as part of a larger thermal stress experiment conducted at the SJINML Biosafety Level-1 aquaria. PAR was monitored daily using using an Apogee MQ-510 underwater quantum meter.

PAM measurements

PAM fluorometry readings were recorded from all coral fragments daily after the final hour of the dark cycle, ensuring full dark adaptation, using a Diving PAM II (Heinz Walz GmbH, Pfullingen, Germany). A saturating pulse (800 ms, default maximal intensity) was applied to determine the maximal fluorescence ($F_m$). The fibre optic probe was positioned using a distance clip to maintain a constant distance and 60° angle between probe and coral sample across measurements. The maximum quantum yield of PSII ($F_v/F_m$) was calculated as ($F_m$-$F_0$)/$F_m$ where $F_0$ is the minimal fluorescence of the dark-adapted sample, measured under the weak measuring light before the saturating pulse.

**Supplementary Note 1**

Essential genes on chromid pCJH

Three essential genes were found on pCJH: *map*, *fabI* and *dnaK*, encoding methionine aminopeptidase (MetAP), enoyl-[acyl-carrier-protein] reductase (FabI) and Hsp70 chaperone, respectively. MetAP and FabI have 50.4% and 42.4% amino acid identity to their characterized homologues in *Escherichia coli* [4, 5] and 38.2% and 44.9% amino acid identity to their respective homologues on SCP18 chromosome. The truncated chromid DnaK lacking a substrate-binding domain is highly divergent from the chromosomal DnaK (23.3% identity) which shares 67.7% amino acid identity to the characterized *E. coli* homologue [6].

Genetic features on pCJH

pCJH encodes functions that could support *Pseudovibrio* symbiosis and persistence within the coral host-associated environment (Fig. 2E). The chromid carries genes associated with amino acid, carbohydrate, lipid and cofactor/vitamin metabolism, together with 51 predicted nutrient and metabolite transport proteins, including 5 complete TRAP-type transporters. Although the core motility and chemotaxis machinery is chromosomally encoded, pCJH expands the sensory repertoire with 3 additional methyl-accepting chemotaxis proteins (MCPs) and a PAS-domain sensory protein potentially related to an aerotaxis receptor, complementing the 15 chromosomal MCPs. Healthy coral polyps become hyperoxic under light, whereas bleached polyps remain hypoxic due to the loss of photosymbionts [7, 8]. The additional aerotaxis capacity of pCJH may facilitate chemotaxis of chromid-containing symbionts towards healthy corals or their positioning within oxygen-rich microenvironments, while additional MCPs could broaden responses to host-derived compounds such as polysaccharides and DMSP [9]. Notably, pCJH also encodes a multidomain c-di-GMP signalling protein, potentially integrating environmental cues to c-di-GMP-dependent regulation of motility and surface attachment.

The expanded sensory repertoire is accompanied by functions that could influence how pCJH-bearing cells interact with the host surface. The chromid encodes a CdiA/FhaB-like two-partner secretion protein and an autotransporter, whose domain architecture is further explored in Fig. 3A, as well as components of capsular polysaccharide biosynthesis and export. Although capsular polysaccharides can be associated with virulence, an analogous system in rhizobia is required for successful attachment to the host [10]. The chromid also encodes RTX-associated extracellular proteins, type I and type III secretion components, although their roles in host interaction remain to be established. pCJH further carries a large NRPS gene cluster homologous to the experimentally characterized *P.*

*brasiliensis ppp* locus, whose products, pseudovibriamide A and B, modulate motility, biofilm formation and overall gene expression [11]. Genes that could help *Pseudovibrio* buffer the coral holobiont against reactive chemical stress [12, 13] are also found on the chromid. These include formaldehyde and methylglyoxal detoxification genes, and peroxidases. Photosynthetic cells generate more methylglyoxal under stress as a Calvin cycle byproduct [14], making its detoxification potentially relevant within the coral holobiont. The chromid also encodes a glutathione-dependent detoxification system, consistent with the proposed role of bacterial ROS-scavenging functions in mitigating oxidative stress within coral holobionts [12, 13, 15], and two other genes involved in nitric oxide detoxification. Furthermore, glycine betaine is abundant in coral tissues, and its metabolism is widespread across coral holobionts, where it contributes to osmoprotection and nitrogen cycling [16]. Although candidate genes for glycine betaine uptake, and downstream sarcosine metabolism are on the chromosome, the chromid encodes a key gene, dimethylglycine dehydrogenase, linking pCJH to glycine-betaine turnover. Finally, two transposases flank a region containing numerous hypothetical proteins, consistent with an evolutionarily dynamic accessory region. Of the 388 genes on pCJH, 87 (22.4%) were annotated as hypothetical, compared with 2,032 of 4,742 (42.9%) chromosomal genes. Thus, the chromid is enriched in genes with assignable functions that expand *Pseudovibrio*'s potential for environmental sensing, host attachment, secondary metabolism and stress tolerance, which could collectively support its symbiosis with the coral holobiont.

**Supplementary Note 2**

<u>Effector-immunity region of T6SS-2</u>

The variable region downstream of *vgrG* in T6SS-2 (Fig. 6F) distinguishes this locus from otherwise similar *Pseudovibrio* T6SS loci. It encodes a polymorphic nuclease effector flanked by putative cognate adapter and immunity proteins, followed by a suspected orphan immunity protein. A homolog of this orphan immunity protein lies downstream of a T6SS-associated endonuclease effector (WP_063296246) in *Pseudovibrio* sp. Ad37, and presumably confers resistance against that effector.

**Supplementary Tables**

**Table S1**: *Pseudovibrio* strains isolated from *Pachyseris speciosa*

| Strain | Isolation source | Colony description | Isolation temp (°C) | Genome size (bp) | No. of CDS | GC content (%) | No. of contigs | Assembly accession number | SRA accession number |
|---|---|---|---|---|---|---|---|---|---|
| SCP18 | Coral macerate | Brown circular colonies | 28 | 5,657,728 | 5,059 | 52.51 | 35 | JBYMDI000000000 | SRR38744195 |
| SCP19 | Coral macerate | Brown circular colonies | 33 | 5,167,238 | 4,671 | 52.48 | 33 | JBDZYJ010000000 | SRR28745652 |
| SCP20 | Coral macerate | Brown circular colonies | 33 | 5,166,901 | 4,669 | 52.48 | 31 | JBYMDH000000000 | SRR38744194 |
| SCP21 | Coral macerate | Brown circular colonies | 33 | 5,657,134 | 5,058 | 52.50 | 37 | JBYMDG000000000 | SRR38744193 |
| SCP23 | Coral macerate | Brown circular colonies | 33 | 5,167,353 | 4,667 | 52.49 | 36 | JBYMDF000000000 | SRR38744192 |
| SCP24 | Coral macerate | Brown circular colonies | 28 | 5,657,236 | 5,061 | 52.51 | 38 | JBYMDE000000000 | SRR38744191 |
| SCP26 | Coral macerate | Brown circular colonies | 33 | 5,167,503 | 4,671 | 52.48 | 36 | JBYMDD000000000 | SRR38744190 |
| SCP27 | Coral macerate | Brown circular colonies | 28 | 5,657,021 | 5,058 | 52.51 | 36 | JBYMDC000000000 | SRR38744189 |
| SCP28 | Coral macerate | Brown circular colonies | 28 | 5,657,087 | 5,058 | 52.51 | 33 | JBYMDB000000000 | SRR38744188 |
| SCP29 | Coral macerate | Brown circular colonies | 23 | 5,656,907 | 5,058 | 52.51 | 35 | JBYMDA000000000 | SRR38744187 |
| SCP30 | Coral macerate | Brown circular colonies | 23 | 5,657,363 | 5,058 | 52.51 | 39 | JBYMCZ000000000 | SRR38744186 |

CDS – coding sequence

**Table S2**: *Pseudovibrio* type strains used in this study

| Strain | Genbank assembly number |
|---|---|
| *Pseudovibrio ascidiaceicola* F423 | GCA_900114245 |
| *Pseudovibrio axinellae* Ad2 | GCA_900110875 |
| *Pseudovibrio brasiliensis* Ab134 | GCA_018282095 |
| *Pseudovibrio denitrificans* DN34 | GCA_900116575 |
| *Pseudovibrio exalbescens* LA33B | GCA_000422785 |
| *Pseudovibrio flavus* RKSG542 | GCA_009711345 |
| *Pseudovibrio japonicus* WSF2 | GCA_014651595 |
| *Pseudovibrio* sp. FO-BEG1 | GCA_000236645 |

**Table S3**: Antibacterial screen of *Pseudovibrio* isolates against *Vibrio* species

| ***Pseudovibrio* sp. strain** | Zone of inhibition (ZOI) in mm[1] | | | | |
|---|---|---|---|---|---|
| | ***Vibrio* sp. SCP2** | ***V. coralliilyticus* SCP4** | ***V. coralliilyticus* YB1** | ***V. harveyi* SCP5** | ***V. harveyi* 384** |
| **SCP18** | 0.5 ± 0 | < 0.5 ± 0 | < 0.5 ± 0 | 2 ± 0 | 2 ± 0 |
| **SCP19** | < 0.5 ± 0 | < 0.5 ± 0 | < 0.5 ± 0 | 2 ± 0 | 2 ± 0 |
| **SCP20** | < 0.5 ± 0 | < 0.5 ± 0 | < 0.5 ± 0 | 2.5 ± 0 | 2 ± 0 |
| **SCP21** | 1 ± 0 | < 0.5 ± 0 | 0 ± 0 | 2.5 ± 0 | 2 ± 0 |
| **SCP23** | 2 ± 0 | < 0.5 ± 0 | 0.5 ± 0 | 3 ± 0 | 2 ± 0 |
| **SCP24** | 0.5 ± 0 | < 0.5 ± 0 | 1 ± 0 | 3 ± 0 | 3 ± 0 |
| **SCP26** | 0 ± 0 | < 0.5 ± 0 | 1 ± 0 | 3 ± 0 | 2 ± 0 |
| **SCP27** | 0 ± 0 | < 0.5 ± 0 | < 0.5 ± 0 | 3 ± 0 | 2 ± 0 |
| **SCP28** | < 0.5 ± 0 | 0.5 ± 0 | 1 ± 0 | 3 ± 0 | 2.5 ± 0 |
| **SCP29** | < 0.5 ± 0 | < 0.5 ± 0 | 1 ± 0 | 3 ± 0 | 2.5 ± 0 |
| **SCP30** | 0.5 ± 0 | < 0.5 ± 0 | 0 ± 0 | 2 ± 0 | 1.5 ± 0 |

[1] Antibacterial screening was performed in triplicate (n = 3)

### Supplementary figures

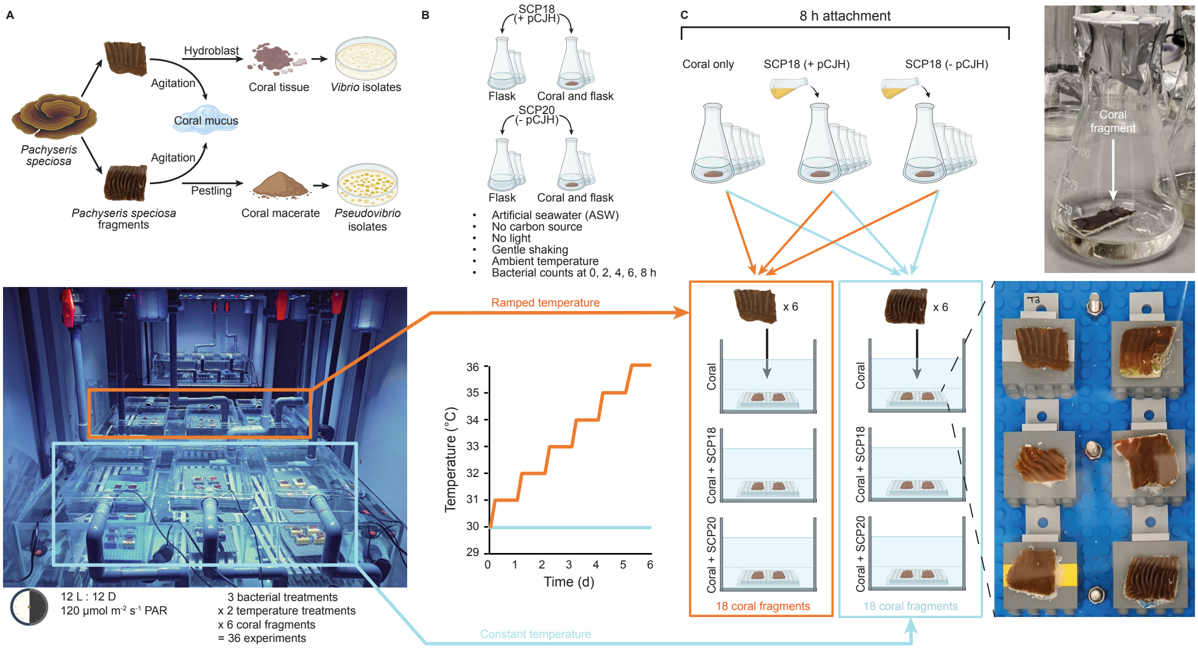


**Figure S1: Experimental setup of bacterial isolation, host colonisation assays, and coral temperature stress experiments.** (A) *Vibrio* and *Pseudovibrio* isolates were obtained from *Pachyseris speciosa* coral tissue and coral macerate, respectively, after agitation to dissociate surface mucus from the coral fragment. (B) Pseudovibrio sp. with chromid (SCP18, + pCJH) and Pseudovibrio sp. without chromid (SCP20, - pCJH) were each inoculated into flasks with or without coral fragment, and bacterial attachment to coral is quantified every 2 h for 8 h via bacterial colony forming unit (CFU) count. (C) SCP18 and SCP20 were each inoculated into flasks containing a coral fragment, alongside a coral-only control. The photo in the top right panel shows an example of coral fragment in its flask. Flasks were incubated for 8 h to allow bacterial attachment to the coral fragments. Six bacteria-treated coral fragments were then mounted on a board (bottom right panel) and placed into a tank held at either constant or ramped temperature (tanks set up shown in bottom left panel). Temperature in the ramped tanks was raised by 1 °C every 24 h. Coral fragments were maintained under these respective conditions for 6 days, yielding a total of 36 experiments in which PSII maximum quantum efficiency ($F_v/F_m$) was measured every 24 h.

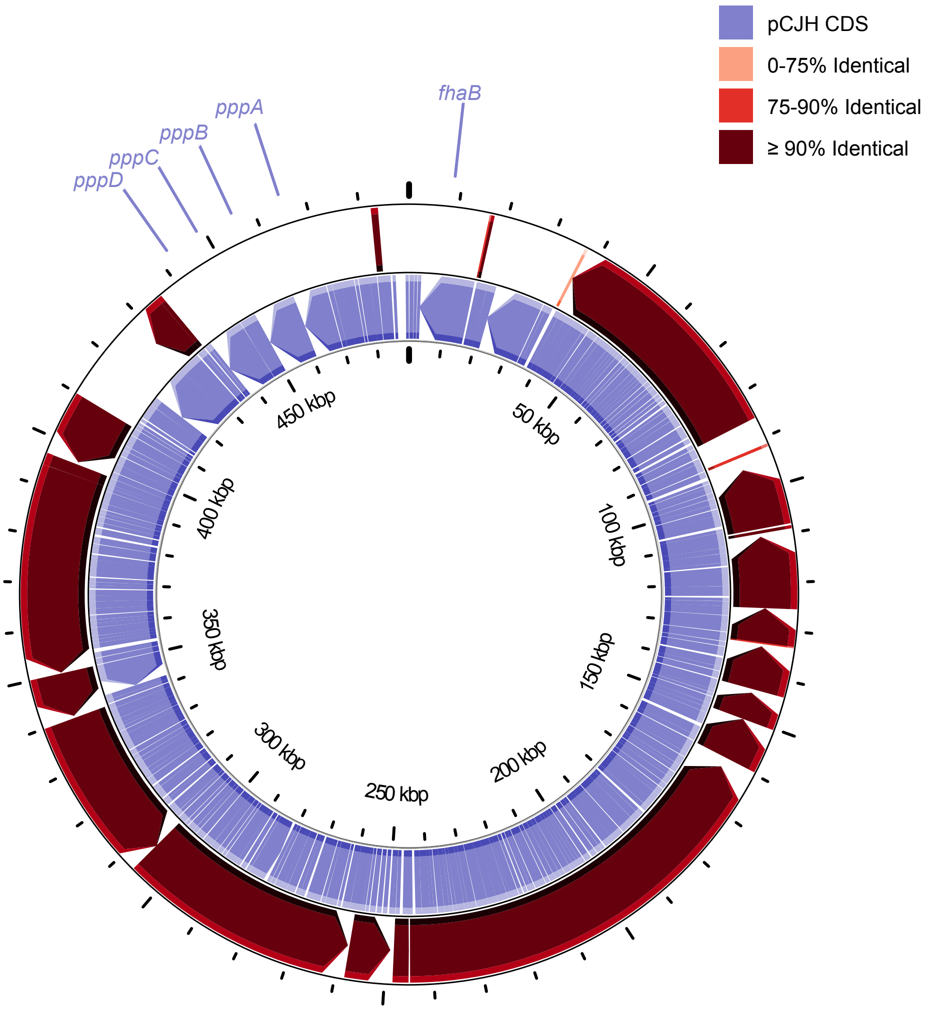


**Figure S2: BLAST atlas comparing the *Pseudovibrio* sp. FO-BEG1 chromid against *Pseudovibrio* sp. pCJH chromid.** The inner track shows the coding domain sequences (CDSs) of pCJH, and the outer track shows BLASTn hits of FO-BEG1 mapped against pCJH. No homologs of the pseudovibriamide (*ppp*) biosynthetic operon or the *cdiA*-like *fhaB* gene were detected on the FO-BEG1 chromid.

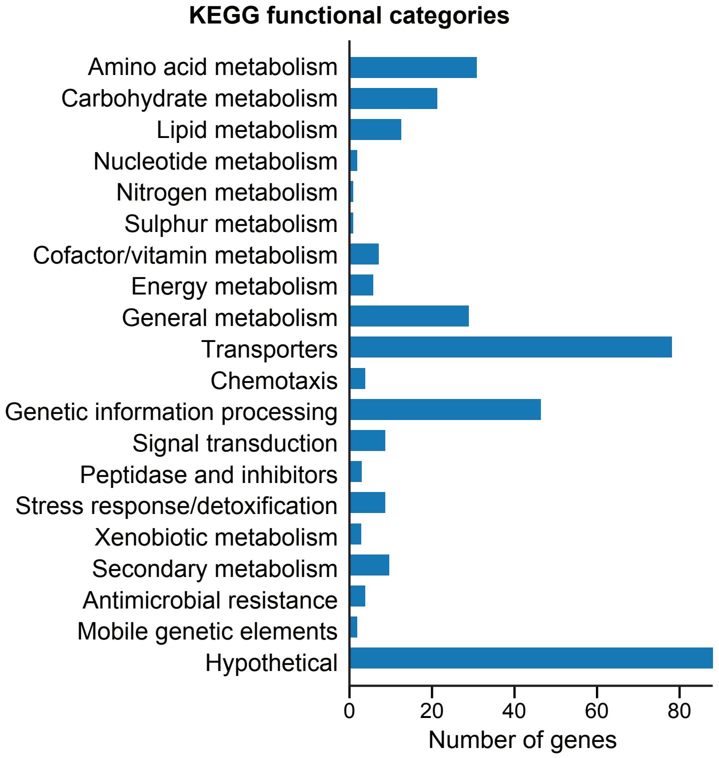


**Figure S3: KEGG functional categories on pCJH.** The 388 genes carried on pCJH were assigned to KEGG or GO functional categories by manual curation, cross-referencing annotations from eggNOG-mapper, Prokka, and BLASTp. Of these, 87 genes (22.4%) were categorized as hypothetical, as they were annotated as hypothetical proteins across all three methods.

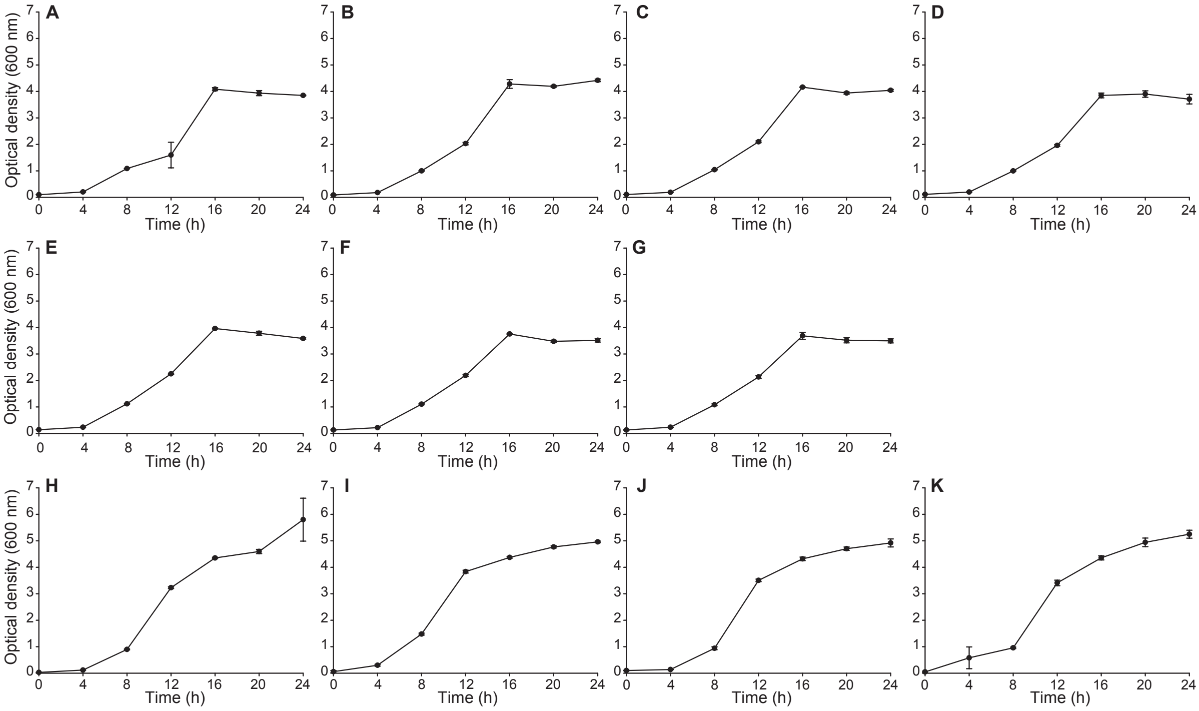


**Figure S4: Growth curves of *Pseudovibrio* sp. isolates.** *Pseudovibrio* sp. SCP18 (A), SCP21 (B), SCP24 (C), SCP27 (D), SCP28 (E), SCP29 (F) and SCP30 (G) harbour the chromid pCJH while *Pseudovibrio* sp. SCP19 (H), SCP20 (I), SCP23 (J) and SCP26 (K) do not.

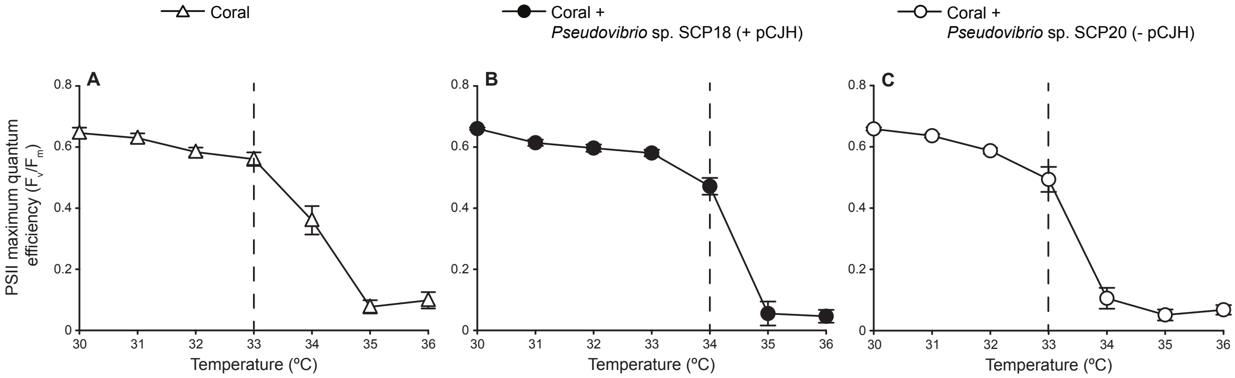


**Figure S5: Individual PSII maximum quantum efficiency ($F_v/F_m$) curves of *P. speciosa* inoculated with SCP18 (+ pCJH) and SCP20 (- pCJH).** Tipping points, defined as the last temperature point before $F_v/F_m$ dropped significantly at the following temperature point, are indicated by a dashed line.

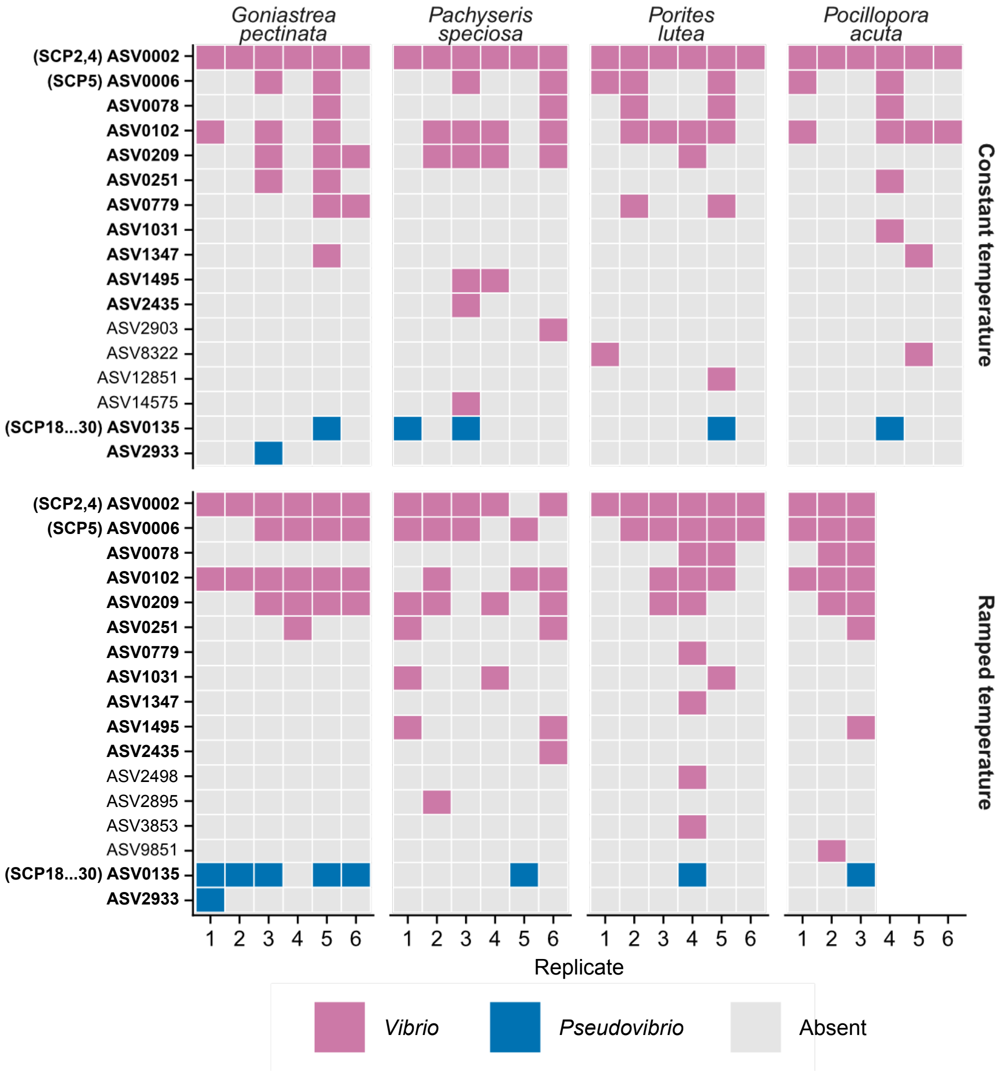


**Figure S6: Presence of amplicon sequence variants (ASVs) in constant and ramped temperature treatments.** ASVs highlighted in bold are present in both experimental conditions. Each column represents individual coral fragments as replicates.
